# The Effects of Hypertension on Signaling Dynamics in Rare Renal Cell Types

**DOI:** 10.64898/2026.01.23.701352

**Authors:** Justin G. McDermott, Bethany L. Goodlett, Shobana Navaneethabalakrishnan, Joseph M. Rutkowski, Brett M. Mitchell

**Author notes:** Correspondence to: Brett M. Mitchell, PhD, FAHA, Address: 8447 John Sharp Parkway, Bryan, TX 77807.

## Abstract

Hypertension (HTN) is the most prevalent risk factor for severe cardiovascular disease and can cause major renal damage, inflammation, and immune cell accumulation. Lymphatic endothelial cells (LECs) are involved in the removal of pro-inflammatory immune cells and cytokines and kidney-specific augmentation of lymphangiogenesis can prevent or reduce HTN. In our previous paper, we performed single-cell RNA sequencing (scRNAseq) on CD31+/podoplanin+ renal cells from mice that underwent angiotensin II-induced (A2HTN) or salt sensitive (SSHTN) models of HTN (and their respective controls) and identified populations of LECs, myeloid immune cells (MICs), and a novel multipotent population we dubbed support cells (SCs). Using NicheNet, we compared baseline signaling between these three cell types in control samples and differences in signaling between control and HTN samples in both LECs and SCs. Ligands with high regulatory potential were identified for all three cell types, with *Tgfb1* having the strongest and most consistent activity across all cell types. When comparing control and HTN samples in both LECs and SCs, HTN samples consistently had a larger number of downstream targets enriched and targets that were enriched in HTN samples also corresponded to significantly increased differentially expressed genes (p<0.01) as reported previously. Significant GO terms (p<0.01) were identified from targets and showed a shift in HTN samples away from homeostatic processes and toward growth and proliferation in LECs and translation and metabolism in SCs. Validation and manipulation of the ligand-receptor-target links identified here may provide novel approaches to reduce renal inflammation and immune cell activation.

**Graphical Abstract:** 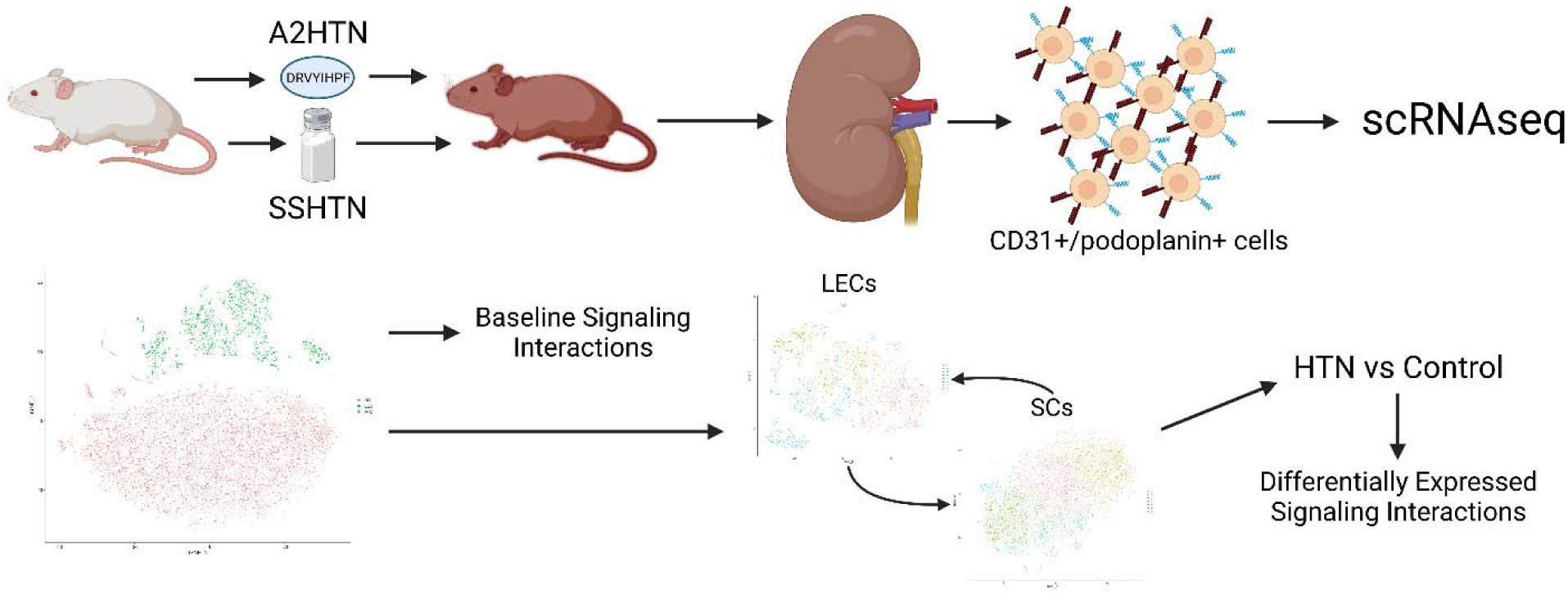

Created with BioRender.com

## Introduction

Hypertension (HTN) is the most common cardiovascular risk factor and a global concern, with its prevalence doubling in the past 30 years.^1, 2^ Despite this, current treatments are ineffective at blood pressure (BP) control in a majority of HTN patients and this leads to systemic organ damage, particularly in the kidneys. In HTN, the kidneys become damaged and inflamed, leading to an accumulation of pro-inflammatory immune cells which further exacerbate the initial issues.^3^

Lymphatic endothelial cells (LECs) are able to remove immune cells and the cytokines they produce from an area in order to reduce inflammation and our previous work has shown that augmenting lymphangiogenesis in the kidneys of mice with HTN can prevent or reduce HTN.^4–8^ To better understand how renal LECs respond to HTN, we utilized single-cell RNA sequencing (scRNAseq) of CD31+/podoplanin+ renal cells from mice that underwent the angiotensin II-induced and salt sensitive models of HTN (A2HTN and SSHTN, respectively).^9–11^ In addition to LECs, we identified a population of myeloid immune cells (MICs) and a novel multipotent population that we call support cells (SCs).^10, 11^

The kidneys consist of an extensive range of cell types which are tightly regulated during development and into adulthood by genetic programs and autocrine and paracrine signaling.^12, 13^ Signaling in renal cell types can be drastically altered by inflammatory conditions, but the specific effects of HTN are unknown.^14^ Cellular crosstalk underlies the constant maintenance of each cell’s identity and function while significant alterations in signaling can result from damage and inflammation, particularly in LECs and stem cells.^15^ To better understand the changes that LECs and SCs experience in HTN, we utilized NicheNet to evaluate the signaling present in each cell type in our HTN models and their controls, as well as between LECs, SCs, and MICs at baseline.

## Methods

Murine hypertension models, CD31+/podoplanin+ enrichment, library preparation and sequencing, quality control, and cell type determination were performed as previously described.^10, 11^ Raw and processed sequencing data are available through NCBI GEO (GSE236410). To evaluate signaling in control samples between LECs, SCs, and MICs and determine if HTN led to differential expression (DE) of signaling to LECs and SCs, NicheNet (nichenetr v1.1.1) was used with default/recommended parameters (except where described otherwise) and an adjusted p-value cutoff of <0.01.^16^ In the control samples, LECs, SCs, and MICs were each evaluated as the receiver cell type and we limited analysis to genes with a minimum log fold change cutoff of 0.25 and ligand-targets pairs with regulatory potential scores ≥0.5. When evaluating for DE signaling, LECs and SCs were each treated as the receiver cell type and we used a log fold change threshold of 0.3, evaluated the top 250 targets, and gave preference to experimentally established (rather than predicted) ligand-receptor-target links. Gene ontology (GO) terms for DE targets for each group were examined using LAGO (https://go.princeton.edu/cgi-bin/LAGO) for all GO aspects with a p-value cutoff of 0.01 and Bonferroni correction applied.^17^

## Results

### Signaling in Normotensive Samples

We chose to first determine what, if any, signaling between LECs, SCs, and MICs was active under baseline conditions in our normotensive control (Ctrl) samples. Ligands that were strongly expressed by only one of the three cell types were determined (Figure 1A). Ligand genes expressed by any cell type were compared against receptor expression in each cell type using a list of known interactions (Figure 1B-1D). These interactions do not distinguish between direct and indirect signaling, though direct signaling does yield a stronger interaction potential score, as seen in Figure 1B. VEGF-A can bind VEGFR1 (*Flt1*) and VEGFR2 (*Kdr*) directly, but not VEGFR3 (*Flt4*).

**Figure 1.**
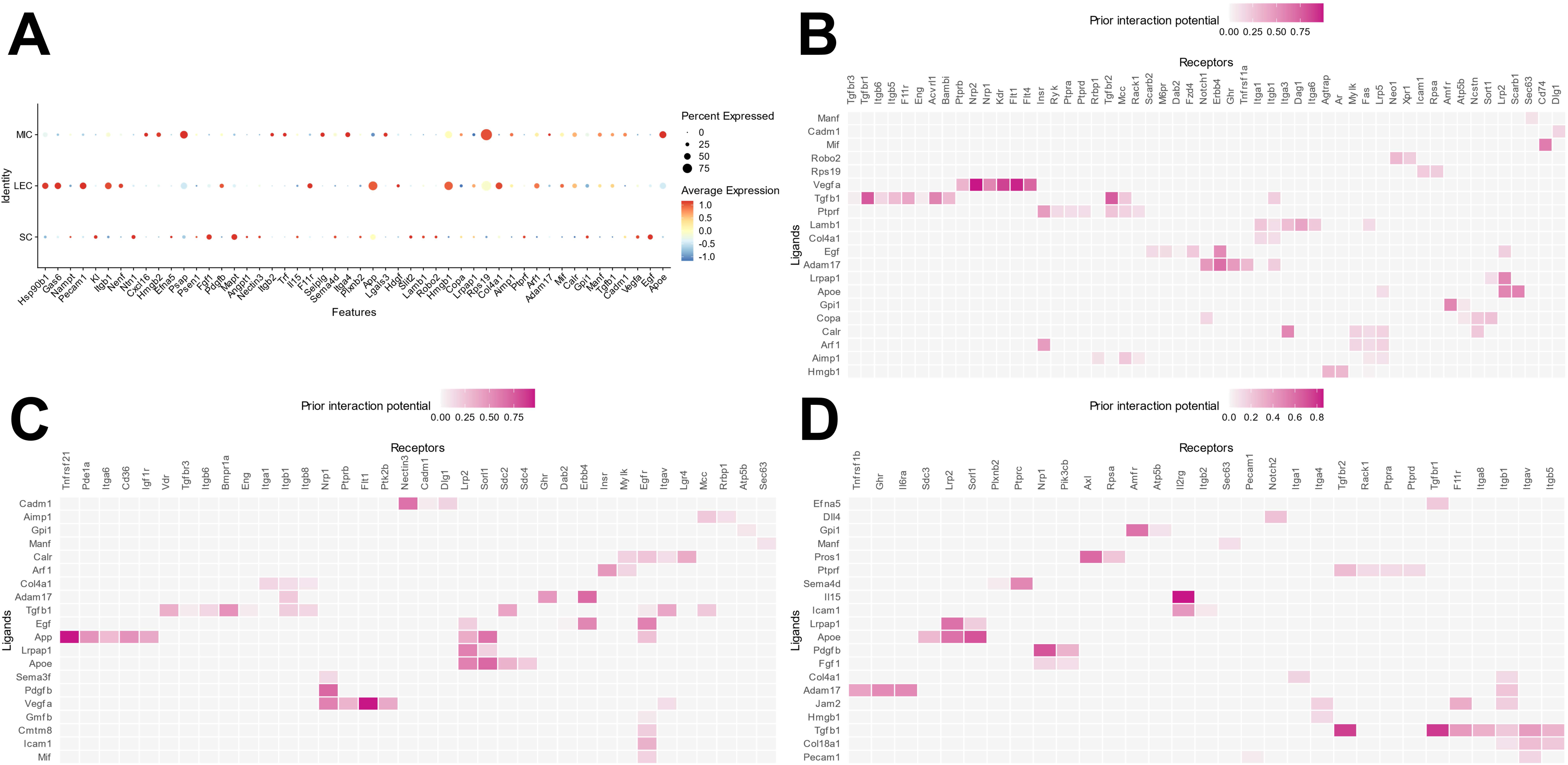
Dot plot of expression for the most highly expressed ligands by cell type (A) and ligand-receptor interaction matrices by cell type (B – LECs, C – SCs, D – MICs).

However, VEGF-A activation on other cell types can induce the production of VEGF-C or VEGF-D, which then bind VEGFR3, and this indirect approach leads to a lower score.^18^ The most active putative ligands for each cell type are *Tgfb1*, *Vegfa*, and *Adam17* (LECs), *App*, *Vegfa*, *Apoe*, and *Tgfb1* (SCs), and *Tgfb1*, *Adam17*, *Apoe*, and *Il15* (MICs).

Based on the list of ligand-receptor interactions for each cell type, downstream target activation by each ligand-receptor pair was also evaluated. Co-regulation scores using the expression of downstream targets in each cell type and the expression of ligands with the highest activity produced by the other cell types were generated and are shown in the right-most plots in Figure 2. Some of the major regulatory ligands for each cell type include *Mif*, *Calr*, *Tgfb1*, *Cadm1*, and *Egf* for LECs (Figure 2A), *App*, *Mif*, *Tgfb1*, *Cadm1*, *Calr*, and *Egf* for SCs (Figure 2B), and *Tgfb1*, *Apoe*, *Adam17*, *Il15*, and *Hmgb1* for MICs (Figure 2C). *Tgfb1* in particular stands out among the other ligands due to its widespread effects across each cell type. Beyond its role as a major driver of downstream target expression through multiple receptors in all cell types, it appears to be indirectly inducing increases in its own expression through intermediates like *Calr* and *Hmgb1*. *Tgfb1* is known to have many regulatory properties during development and adulthood and it may be helping to maintain homeostasis in these cell types through its ability to dampen proliferative and pro-inflammatory pathways.^19, 20^ It should also be noted from these figures that LECs and MICs produce the majority of active ligands while SCs have the largest number of regulated downstream targets and the highest average regulatory potential for targets, which may indicate that they are more susceptible to alterations in expression due to external factors.

**Figure 2.**
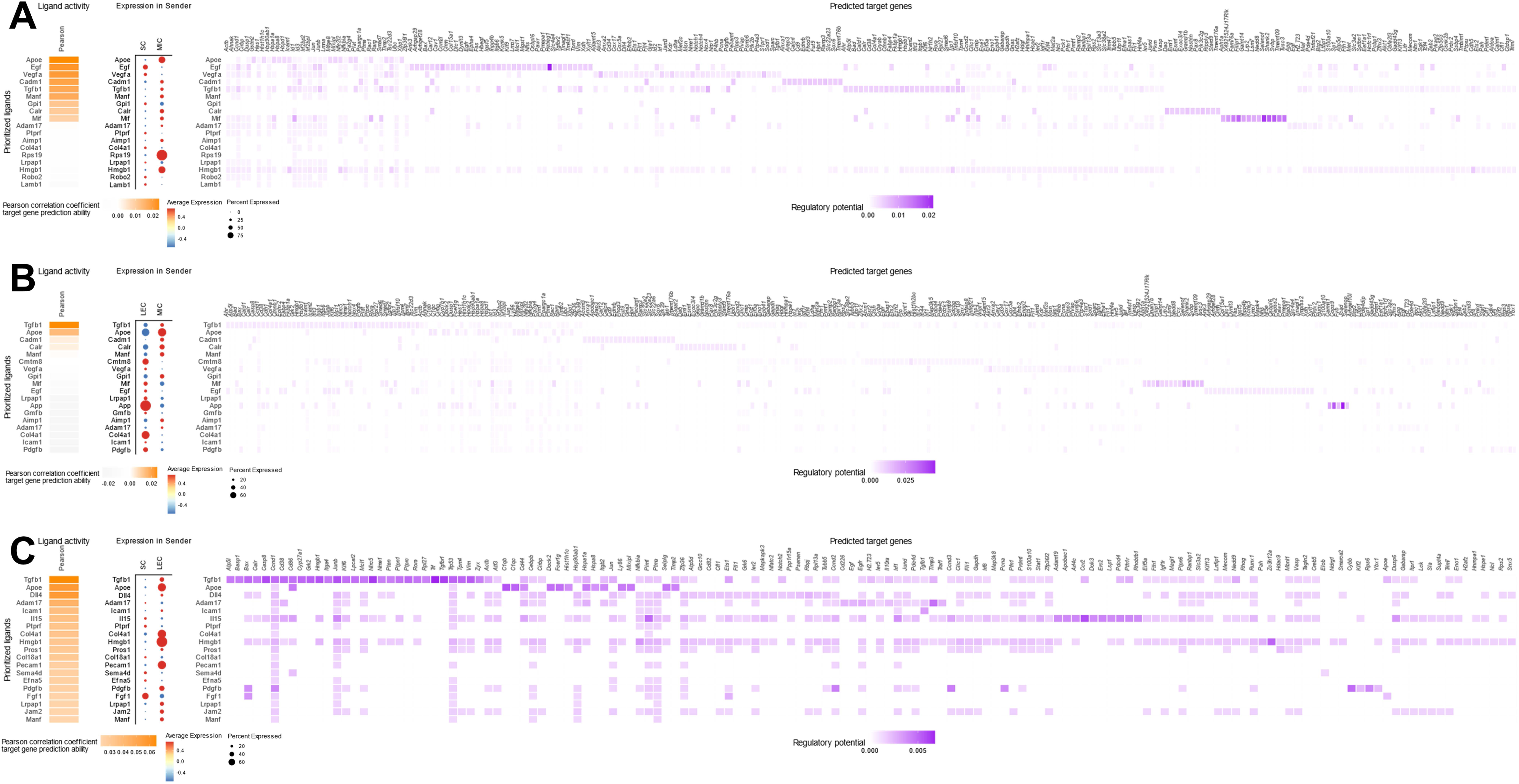
Ligand activity, expression, and regulatory potential plots with each cell type acting as the receiver (A – LECs, B – SCs, C – MICs).

### Signaling Alterations in LECs in HTN

To evaluate the changes in signaling resulting from HTN, we compared our Ctrl and HTN samples with LECs or SCs as the recipient cell type. With LECs as the signaling recipients, we identified ligand-receptor pairs that were upregulated in either Ctrl or HTN samples (Figure 3). Roughly half of the ligands enriched in Ctrl samples (labeled as “Ctrl_niche”) corresponded to increases in their paired receptors (Figure 3A), whereas the majority of ligands enriched in HTN (labeled as “HTN_niche”) had concomitant increases in their paired receptors (Figure 3B). In A2HTN and SSHTN-only comparisons to Ctrl samples, a similar trend emerged with more consistent upregulation of both ligands and receptors from each pair in HTN samples (Figures S1&S2).

**Figure 3.**
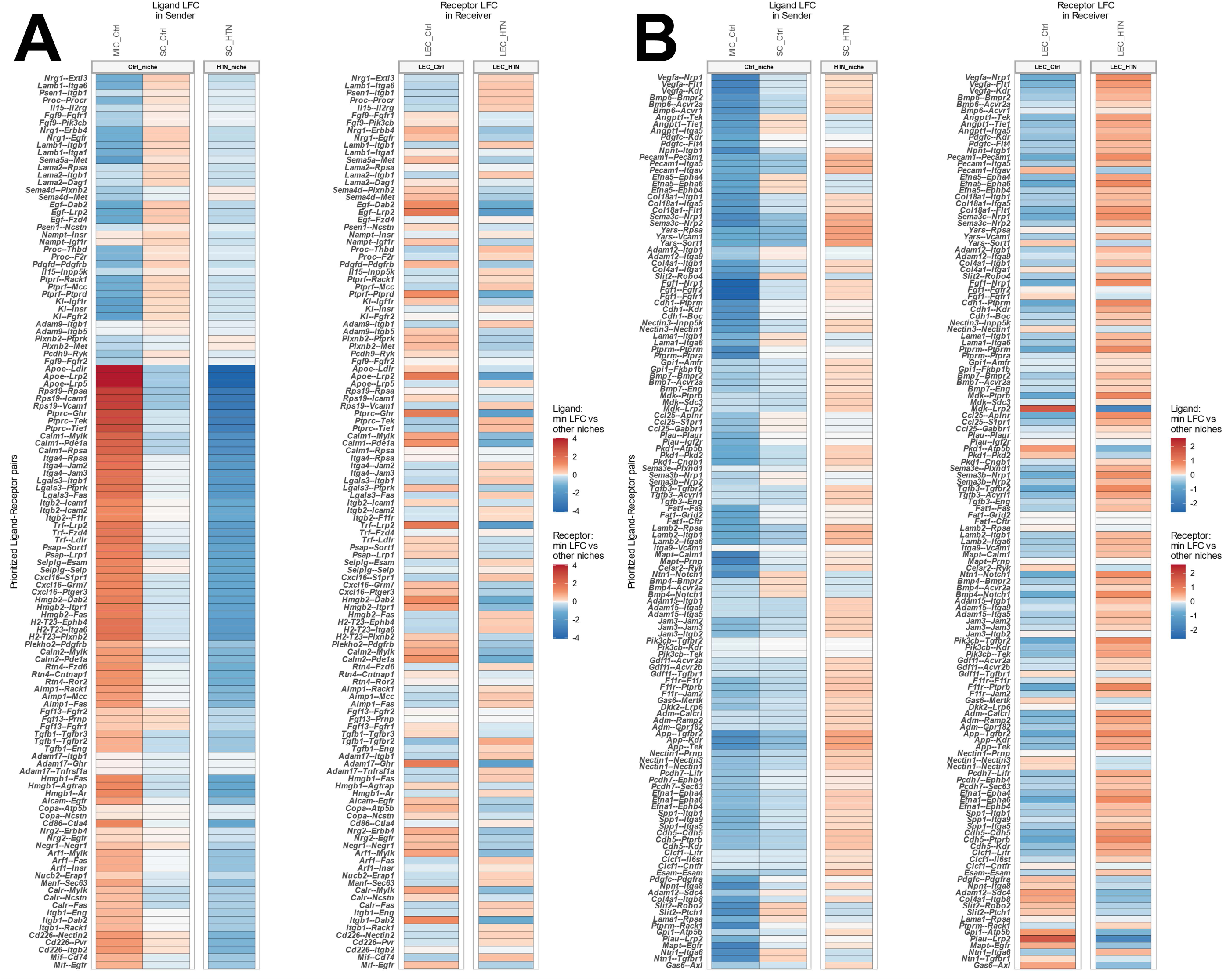
Differential expression of ligands and receptors with LECs as the receiver cell type and comparing the Ctrl group (A) versus the merged HTN group (B). Ligands and receptors are shown in pairs (Ligand--Receptor) and changes in expression are visualized through log fold change (LFC) values.

Summary plots of the ligands and receptors for each group and condition can be found in Figure S3.

Ligand-receptor links with strong ligand expression in either Ctrl or HTN samples were then compared to downstream target expression to assess their unscaled and scaled activity, as well as their impact on target genes that were altered in each condition (Figure 4). There were far fewer targets active in LECs in the Ctrl samples (Figure 4A) compared to the targets active in the HTN samples (Figure 4B), reinforcing the idea that paracrine signaling between these cell types is involved in pro-inflammatory changes in LECs. Similar comparisons with A2HTN and SSHTN versus Ctrl are shown in Figures S4&S5.

**Figure 4.**
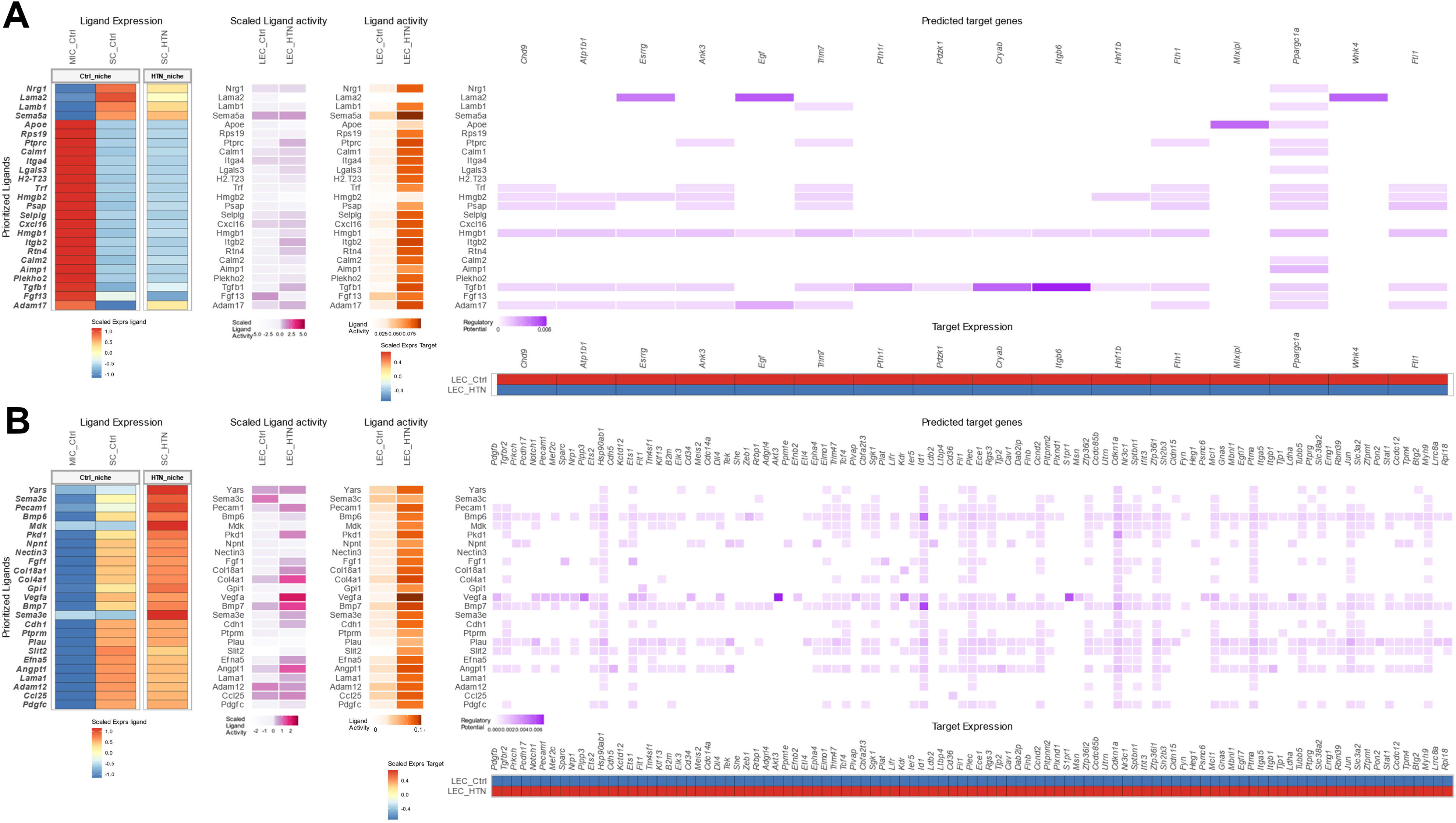
Ligand expression, activity, and regulatory potential plots with LECs as the receiver cell type and showing DE ligands with high activity in the Ctrl group (A) or the merged HTN group (B).

Some of the major ligand-target links in the Ctrl samples include *Apoe*—*Mlxipl*, *Lama2*—*Esrrg*, and *Tgfb1*—*Itgb6*. Each of these targets were downregulated in HTN (*Mlxipl*, 69% decrease, p<0.0001; *Esrrg*, 212% decrease, p<0.0001; *Itgb6*, 86% decrease, p<0.0001) and have anti-inflammatory or anti-proliferative effects.^21–23^ In HTN, *Bmp6*—*Id1*, *Bmp7*—*Id1*, *Vegfa*—*Akt3*, *Vegfa*—*S1pr1*, and *Vegfa*—*Dll4* were among the most active links and had their target genes significantly upregulated (*Id1*, 41% increase, p<0.0001; *Akt3*, 48% increase, p<0.0001; *S1pr1*, 35% increase, p=0.002; *Dll4*, 50% increase, p=0.0006). Bone morphogenetic proteins (BMPs) are part of the TGFβ superfamily, possess angiogenic effects, and are speculated to be involved in lymphangiogenesis as well.^24^ BMP6 can induce sprouting through VEGFR2, but has also been reported to downregulate VEGFR2 expression through TAZ.^25^ ID1 is vital for embryonic vascular development, involved in angiogenesis and maintenance of endothelial identity, is strongly activated by BMPs, and can even prevent endothelial-to-mesenchymal transition induced by TGFβ2.^26, 27^ S1PR1 and DLL4 can have antagonistic effects on lymphatic sprouting resulting from laminar sheer stress, with DLL4 enhancing VEGFR3 signaling and S1PR1 reducing it, despite both being activated by VEGF-A.^28^

Targets specific to each group were analyzed using LAGO to find enriched GO terms from all GO aspects (Tables S1-S6). The most significant GO terms for targets in the Ctrl samples include homeostatic process, chemical homeostasis, inorganic ion homeostasis, and cellular homeostasis (total significant GO term count of 22). In the HTN samples, the most significant terms include anatomical structure morphogenesis, tube morphogenesis, developmental process, vasculature development, cell differentiation, tube development, and cell population proliferation (total significant GO term count of 547).

### Signaling Alterations in SCs in HTN

We also evaluated the effects of HTN on signaling with SCs as the receiver cell type using the same approach as described above with LECs. Ligands expressed by LECs and MICs with strong preferential expression in Ctrl or HTN samples were identified and compared to the expression of their corresponding receptor(s) in SCs (Figure 5). Similar to the Ctrl-specific ligand-receptor pairs for LECs, expression of some of the receptors in both Ctrl and HTN samples was inconsistent with the expression of their paired ligands, indicating a poor link within the pairs, though this did improve slightly in the HTN samples. This trend also applied to the comparisons between the individual HTN samples and Ctrl samples, though A2HTN had slightly more consistent paired receptor expression than SSHTN (Figures S6&S7). Summary plots for the ligand-receptor pairs evaluated for each group and condition are shown in Figure S8.

**Figure 5.**
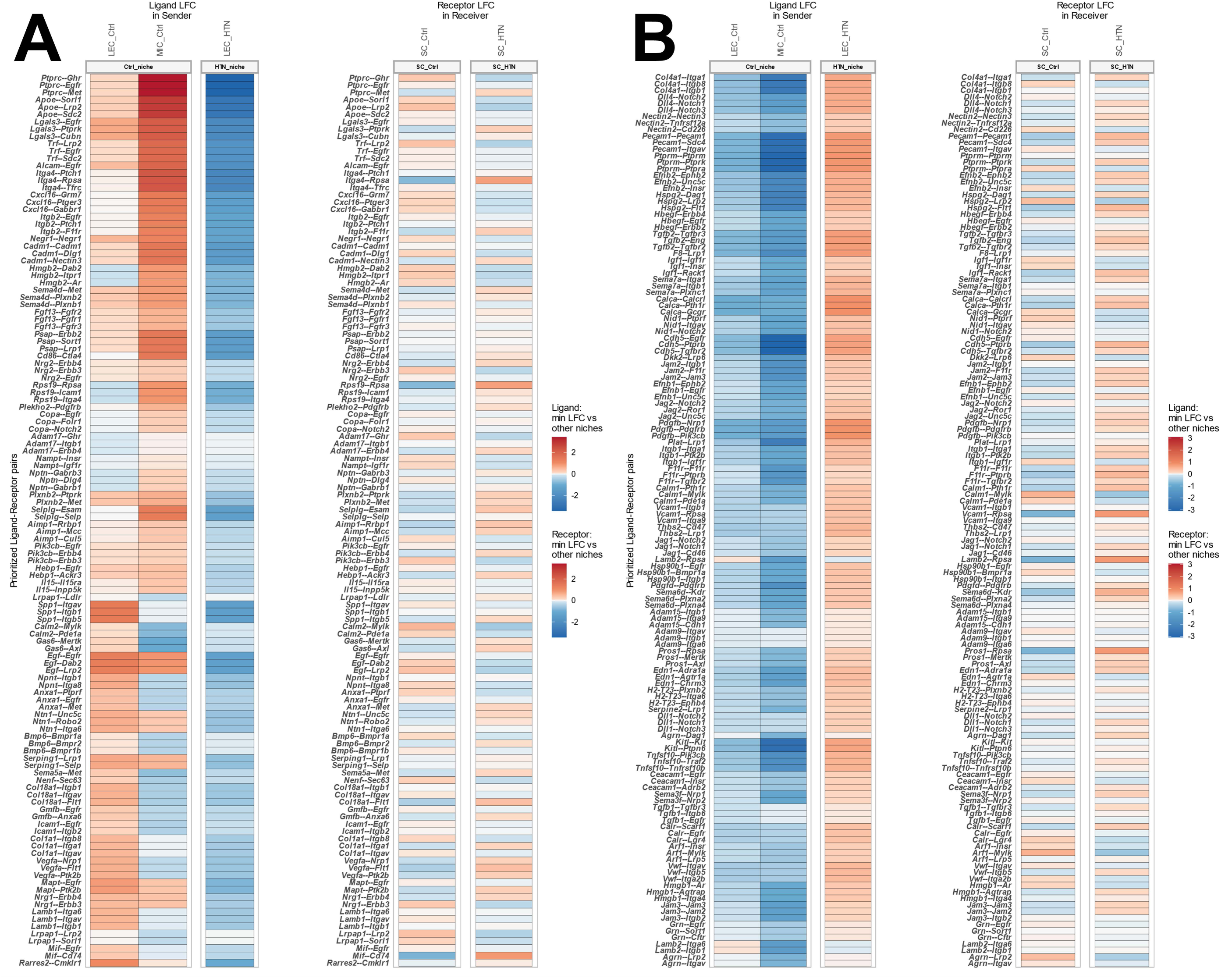
Differential expression of ligands and receptors with SCs as the receiver cell type and comparing the Ctrl group (A) versus the merged HTN group (B). Ligands and receptors are shown in pairs (Ligand--Receptor) and changes in expression are visualized through log fold change (LFC) values.

Ligand-receptor pairs with the highest co-expression and preferential expression in Ctrl or HTN samples were evaluated for downstream target activation (Figure 6). Once again, there were fewer targets active in the Ctrl samples (Figure 6A) compared to the HTN samples (Figure 6B) with a roughly threefold difference between the groups and fewer targets in both Ctrl and HTN samples when compared to their equivalents with LECs as the receiver cell type. Comparisons with the separate HTN samples versus Ctrl samples are present in Figures S9&S10.

**Figure 6.**
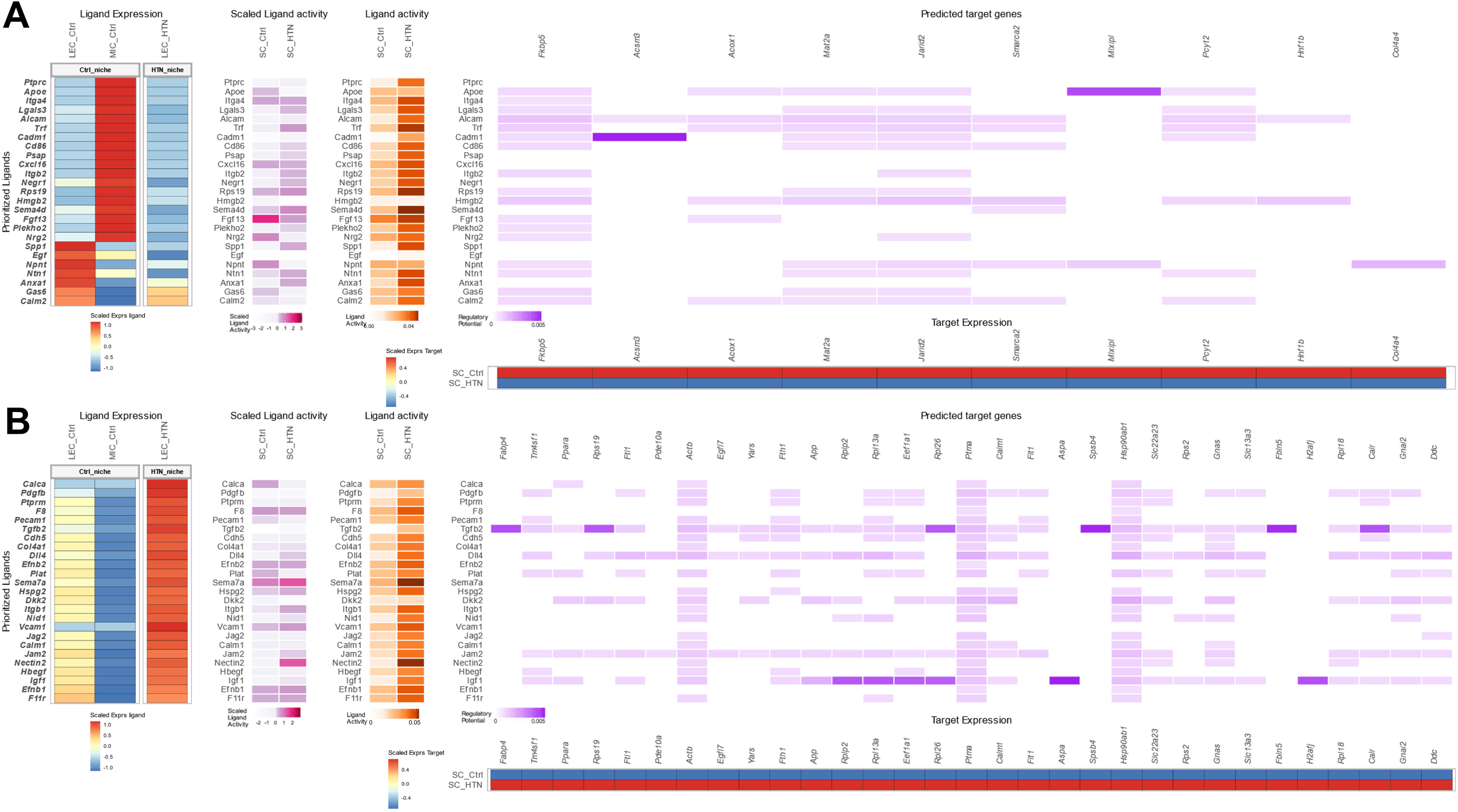
Ligand expression, activity, and regulatory potential plots with LECs as the receiver cell type and showing DE ligands with high activity in the Ctrl group (A) or the merged HTN group (B).

In the Ctrl samples, *Apoe*, *Alcam*, and *Hmgb2* were the most widely active ligands and *Fkbp5* and *Jarid2* were the most consistently activated downstream targets. *Fkbp5* and *Jarid2* are involved in the repression of growth and inflammation and their expression decreased by 39% and 27% respectively (p<0.0001) in HTN samples.^29, 30^ The strongest ligand-target links in these samples were *Cadm1*—*Acsm3* and *Apoe*—*Mlxipl*, with *Acsm3* decreasing by 37% and *Mlxipl* by 23% in HTN samples (p<0.0001). Akin to *Fkbp5* and *Jarid2*, these target also possess anti-proliferative and anti-inflammatory effects.^21, 31^ *Tgfb2*, *Igf1*, *Dll4*, and *Jam2* were the most active ligands in the HTN samples, with *Tgfb2* and *Igf1* having the strongest downstream effects. *Actb*, *Ptma*, and *Hsp90ab1* were the most widely activated targets, were upregulated by 55%, 38%, and 30% respectively (p<0.0001), and are involved in motility and pro-immune and pro-survival pathways.^32–34^ *Tgfb2* was strongly associated with *Fabp4*, *Rps19*, *Rpl26*, *Spsb4*, *Fbln5*, and *Calr* (with increases of 168%, 62%, 41%, 32%, 26%, and 25% in HTN respectively; p<0.0001) while *Igf1* was linked to *Rplp2*, *Rpl13a*, *Eef1a1*, *Rpl26*, *Aspa*, and *H2afj* (with increases of 45%, 42%, 41%, 41%, 33%, and 26% in HTN respectively; p<0.0001). *Fabp4* is upregulated by inflammation and further contributes to pro-inflammatory processes and *Rplp2* is a major component of translational complexes and has pro-survival effects.^35, 36^

Enriched GO terms for targets in SCs from each group were identified by LAGO using the same settings as described above but yielded far fewer terms when compared to targets active in LECs (Tables S7-S12). Targets active in the Ctrl samples only yielded three significant terms—cellular biosynthetic process, organic substance biosynthetic process, and biosynthetic process. Targets from the HTN samples were matched to 62 terms with the top terms including cytosolic ribosome, translation, peptide biosynthetic process, cytoplasmic translation, peptide metabolic process, positive regulation of metabolic process, and regulation of protein metabolic process.

## Discussion

### Intercellular Signaling and Differential Gene Expression

To better contextualize the changes resulting from HTN in both LECs and SCs that were reported previously,^10, 11^ we evaluated the signaling dynamics between these cell types in control and HTN conditions. Despite only having access to LECs and SCs, we were able to identify ligand-receptor-target links with enriched targets that corresponded to significantly altered gene expression in the receiver cell type, with the majority of target genes identified in Figures 4&6 also being significantly upregulated in their respective cell types in each condition. Considering how many different cell types can coexist within relative proximity of each other in the kidneys, the idea that these two cell types in particular can influence one another to such a high degree is notable and suggests that these cell types may even constitute a paracrine signaling niche, especially during inflammation. That said, additional sequencing of other renal cell types from murine HTN models would be immensely beneficial in scrutinizing these interactions and determining if they are actually unique to LECs and SCs.

### Similarities and Distinctions between LECs and SCs in HTN

Despite the potential coordination between LECs and SCs, the downstream targets within each cell type differ widely. With the exceptions of *Mlxipl* and *Hnf1b* in Ctrl samples and *Ptma*, *Hsp90ab1*, *Rpl18*, *Egfl7*, *Flt1*, and *Gnas* in HTN samples, the majority of enriched targets are unique to each cell type. This trend also holds true with the ligands, though each cell type in HTN samples is activated by one of two plasminogen activators (*Plau* in LECs and *Plat* in SCs). These two share an interesting connection in HTN LECs and SCs, with *Fgf1* activating *Plat* in LECs and *Plat* activating *Ptma* and *Hsp90ab1* in SCs, while *Plau* produced by SCs activates *Ptma* and *Hsp90ab1* in LECs. While the two variants both promote the conversion of plasminogen to plasmin, urokinase plasminogen activator (uPA) is generally being bound to its receptor and tissue plasminogen activator (tPA) needs to bind fibrin as a cofactor.^37^ Further work is needed to determine how the distinctions in functionality between uPA and tPA are impacting these cell types in addition to how an increase in plasmin generation, which can have pro-inflammatory effects, is affecting the overall state of damage in kidneys in HTN.^38^

Another major difference between the two cell types is in the number of preferentially activated downstream targets and their corresponding GO terms. LECs simply have more enriched targets in Ctrl and HTN samples than SCs do and this leads to a massive discrepancy in GO term count, with LECs having 547 significant GO terms in HTN samples (compared to 62 in SCs) and 22 terms in Ctrl samples (compared to 3 in SCs). Targets are determined based on their enrichment in one condition over the other and the strength of their association similarly enriched ligand-receptor links, so low target counts do not necessarily indicate a lack of activity, but rather a smaller shift in terms of identifiable linkages. LECs undergo a wide array of changes in inflammation and the number of targets and GO terms reflects this, whereas SCs seem to be more stable in their activity across conditions with lower counts.

### TGFβ Superfamily Signaling Shifts

TGF-β1 is an exceptionally multifaceted cytokine with varying effects on nearly every cell type, though two of its most notable roles are in immunoregulation and stem cell maintenance. In developing immune cells and the early stages of the immune response, it has a proliferative, activating effect, while in the later stages it has an inhibitory effect, suppressing pro-inflammatory cells and reducing cytotoxicity.^39^ Stem cell identity and survival are promoted by and preserved through TGF-β1 signaling and other TGF-β superfamily members are also capable of pushing differentiated cells towards a mesenchymal state.^26, 40^

In the data presented here, we noted that *Tgfb1* is active as a ligand for all cell types at baseline and preferentially active in the Ctrl LEC cells when compared to their HTN counterparts. Instead, in HTN LECs, *Bmp6* and *Bmp7* are active, indicating a shift specific to LECs in inflammatory conditions within the superfamily’s signaling. BMP6 can lead to a decrease in activation and increase in pro-inflammatory cytokine production in macrophages and both BMP6 and BMP7 can negatively impact the growth and production of immunoglobulins by B cells.^39^ While LECs express more immune and stem-associated markers when compared to blood endothelial cells, it is still unclear what exact role(s) these two proteins may play, such as regulating proliferation and production of pro-inflammatory cytokines or augmenting lymphangiogenesis, both at baseline and during inflammation. When comparing Ctrl and HTN SCs, *Tgfb1* didn’t show preferential activity in either group, which may be due to consistent activation needed for the maintenance of their multipotent state, though *Tgfb2* did show preference for activity in HTN. Again, it is unknown exactly what function TGF-β2 has here, as TGF-β1 and TGF-β2 have a great deal of overlap in their roles, though it has been shown that TGF-β2 is a more potent effector of mesenchymal transition in differentiated cells.^26^

## Conclusion

Using NicheNet, we identified signaling activity specific to LECs, SCs, and MICs at baseline conditions and signaling specific to baseline and HTN conditions in both LECs and SCs. LECs in HTN were enriched for a large number of targets and pathways related to growth and proliferation, whereas SCs in HTN had fewer targets with pathways involved in regulation of proliferation and maintenance of stem identity.

Enriched downstream targets in both cell types in each condition largely corresponded to significantly upregulated genes that we identified previously,^10, 11^ and we noted multiple roles of the TGFβ superfamily across cell types and conditions. To better contextualize our findings, additional sequencing of other renal cell types from murine HTN models and validation of ligand-receptor-target links and their effects will be necessary. Manipulation of the signaling axes described here through enhancement or blockade by receptor agonists and antagonists may provide novel approaches to reducing renal inflammation and preventing further renal immune cell accumulation and damage due to HTN.

---

Figure S1 – Differential expression of ligands and receptors with LECs as the receiver cell type and comparing the Ctrl group (A) versus the A2HTN group (B). Ligands and receptors are shown in pairs (Ligand--Receptor) and changes in expression are visualized through log fold change (LFC) values.

Figure S2 – Differential expression of ligands and receptors with LECs as the receiver cell type and comparing the Ctrl group (A) versus the SSHTN group (B). Ligands and receptors are shown in pairs (Ligand--Receptor) and changes in expression are visualized through log fold change (LFC) values.

Figure S3 – Circos plots showing ligands and receptors with LECs as the receiver cell type for the Ctrl group (top row) and HTN groups (bottom row). A and D present ligand-receptor pairs that are DE when comparing the Ctrl group (A) and the merged HTN group (D). B and E present ligand-receptor pairs that are DE when comparing the Ctrl group (B) and the A2HTN group (E). C and F present ligand-receptor pairs that are DE when comparing the Ctrl group (C) and the SSHTN group (F).

Figure S4 – Ligand expression, activity, and regulatory potential plots with LECs as the receiver cell type and showing DE ligands with high activity in the Ctrl group (A) or the A2HTN group (B).

Figure S5 – Ligand expression, activity, and regulatory potential plots with LECs as the receiver cell type and showing DE ligands with high activity in the Ctrl group (A) or the SSHTN group (B).

Figure S6 – Differential expression of ligands and receptors with SCs as the receiver cell type and comparing the Ctrl group (A) versus the A2HTN group (B). Ligands and receptors are shown in pairs (Ligand--Receptor) and changes in expression are visualized through log fold change (LFC) values.

Figure S7 – Differential expression of ligands and receptors with SCs as the receiver cell type and comparing the Ctrl group (A) versus the SSHTN group (B). Ligands and receptors are shown in pairs (Ligand--Receptor) and changes in expression are visualized through log fold change (LFC) values.

Figure S8 – Circos plots showing ligands and receptors with SCs as the receiver cell type for the Ctrl group (top row) and HTN groups (bottom row). A and D present ligand-receptor pairs that are DE when comparing the Ctrl group (A) and the merged HTN group (D). B and E present ligand-receptor pairs that are DE when comparing the Ctrl group (B) and the A2HTN group (E). C and F present ligand-receptor pairs that are DE when comparing the Ctrl group (C) and the SSHTN group (F).

Figure S9 – Ligand expression, activity, and regulatory potential plots with LECs as the receiver cell type and showing DE ligands with high activity in the Ctrl group (A) or the A2HTN group (B).

Figure S10 – Ligand expression, activity, and regulatory potential plots with LECs as the receiver cell type and showing DE ligands with high activity in the Ctrl group (A) or the SSHTN group (B).

## Declaration

### Funding

This work was funded by NIH R01 (DK120493) to B.M. Mitchell and Texas A&M University T3 to J.M. Rutkowski.

## Acknowledgement

N/A

## Author Contributions

Justin G. McDermott-conceptualization, data curation, formal analysis, investigation, methodology, project administration, resources, software, writing-original draft, writing-review and editing; Bethany L. Goodlett-data curation, investigation; Shobana Navaneethabalakrishnan-data curation, investigation; Joseph M. Rutkowski-conceptualization, funding acquisition, project administration; Brett M. Mitchell-conceptualization, funding acquisition, project administration, resources, writing-review and editing

## Conflict of Interest

The authors have nothing to disclose. Consent: N/A

## Ethics

IACUC Animal Use Protocol TAMU #2022-0083

## Data Availability

Raw and processed sequencing data are available through NCBI GEO (GSE236410). All scripts for analysis and their direct results are available upon reasonable request to the corresponding author.

## Notes

https://www.ncbi.nlm.nih.gov/geo/query/acc.cgi?acc=GSE236410

### Competing Interest Statement

The authors have declared no competing interest.

